# Genome-enabled prediction models for black tea (*Camellia sisnesnsis*) quality and drought tolerance traits

**DOI:** 10.1101/850792

**Authors:** Robert. K. Koech, Pelly M. Malebe, Christopher Nyarukowa, Richard Mose, Samson M. Kamunya, Theodor Loots, Zeno Apostolides

## Abstract

**Summary:**

- Genomic selection in tea (*Camellia sinensis*) breeding has the potential to accelerate efficiency of choosing parents with desirable traits at the seedling stage.
- The study evaluated different genome-enabled prediction models for black tea quality and drought tolerance traits in discovery and validation populations. The discovery population comprised of two segregating tea populations (TRFK St. 504 and TRFK St. 524) with 255 F_1_ progenies and 56 individual tea cultivars in validation population genotyped using 1 421 DArTseq markers.
- Two-fold cross-validation was used for training the prediction models in discovery population, and the best prediction models were consequently, fitted to the validation population.
- Of all the four based prediction approaches, putative QTLs (Quantitative Trait Loci) + annotated proteins + KEGG (Kyoto Encyclopaedia of Genes and Genomes) pathway-based prediction approach, showed robustness and usefulness in prediction of phenotypes.
- Extreme Learning Machine model had better prediction ability for catechin, astringency, brightness, briskness, and colour based on putative QTLs + annotated proteins + KEGG pathway approach.
- The percent variables of importance of putatively annotated proteins and KEGG pathways were associated with the phenotypic traits. The findings has for the first time opened up a new avenue for future application of genomic selection in tea breeding.

## Introduction

Tea plant (*Camellia sinensis* (L.) Kuntze) is an important raw material for universally most popular non-alcoholic beverage (Chen *et al.*, 2007). Because tea is an important economic crop, it is grown in tropical and semi-tropical countries around the world including China, India, Kenya and Sri Lanka for production of tea beverages (FAO 2015a). In Kenya, tea production is a major foreign exchange earner, and export commodity among agricultural produces with over 26% of all foreign exchange (Kenya National Bureau of Statistics, 2012; TBK, 2012). Since tea is the most popular beverage globally, the need to develop tea cultivars with optimum potential for black or green tea quality has recently become the single most important breeding objective. However, tea breeding programs have had challenges in breeding tea cultivars with desirable traits, because tea plants are self-incompatible, highly heterozygous, and has a long juvenile period of 4-5 years and about 22-25 years to breed a new cultivar (Chen *et al.*, 2007). In addition, exploiting QTLs for more complex traits such as tea quality and drought tolerance to predict candidates for selection in practical breeding programs has had limited impact because of the small genetic variance accounted for by QTLs owing to the model being used, and many detected QTLs are specific to particular genetic background. Due to the limitations posed by QTLs mapping technique, machine learning methods provide an alternative approach in plant breeding. Machine learning methods have been used in many areas such as genomics, transcriptomics, proteomics, and systems biology (Larranaga *et al.*, 2006; Tarca *et al.*, 2007; Kelchtermans *et al.*, 2014). In animals, machine learning methods have been applied widely in genome research studies which include, genome assembly, genome annotation and gene regulatory network studies, and in prediction of gene function studies (Larranaga *et al.*, 2006; Tarca *et al.*, 2007). Machine learning approaches have been used recently in plants in identification of healthy and non-healthy citrus leaves (ur Rahman *et al.*, 2017), disease resistance in wheat (Ornella *et al.*, 2017), breeding of drought-tolerant maize (Ornella *et al.*, 2012), grain yield in maize and wheat (Shekoofa *et al.*, 2014; Pantazi *et al.*, 2016) and in stress severity studies in soybean (Naik *et al.*, 2017). Also, machine learning has been used in fruit trees (apples) which have a long breeding cycle to select for fruit quality traits (Kumar *et al.*, 2012).

In tea plants, tremendous progress has been made regarding the availability of good genetic resources (tea genome) and linking molecular markers to QTL of complex traits (Hazra *et al.*, 2018). However, the development of new tea cultivars, tea breeders are faced with challenges of choosing among many potential selection criteria. These selection criteria include yield per hectare, cup quality, and tolerance to both abiotic and biotic factors. Besides the yield, the quality of tea is also important from an economic perspective, as quality is an essential factor influencing the market price (Yan, 2007). The price of black tea in the world market depends primarily on the physical appearance of the made tea and the infused leaf which include the aroma and liquor characters (colour, brightness, strength, and briskness) (Bhatia, 1963). The quality of tea, especially, appearance, taste, and smell aspects, is largely determined by the biochemical composition of the fresh tea leaves (Obanda *et al.*, 1997; Dutta *et al.*, 2011). Therefore, this implies that only producers of high-quality black teas are likely to survive in the global tea market.

Drought is a major environmental stress factor that affects the growth and development of tea plants, and the traits associated with drought tolerance are quantitative in nature. Furthermore, climate change and population growth have put more pressure on traditional tea growing regions in Kenya with less land available to conduct field trials on newly developed tea cultivars (FAO, 2015b). This has contributed to tea farming and production being moved to marginal areas which are unsuitable for tea farming and production (FAO, 2015b). Therefore, the objective of tea breeding programs is to develop tea cultivars for production of high-quality black tea and tea cultivars that can adapt to changing environmental conditions such as drought stress. This can be accelerated by applying new and novel techniques such as machine learning methodologies which are more robust, accurate, efficient, and capable of evaluating a much wider set of variables within a short period of time (Crossa *et al.*, 2017). Machine learning methodologies can also help in the identification of genomic regions that are of agronomic value by facilitating functional annotation of genomes and enable real-time high-throughput phenotyping of agronomic traits (Hu *et al.*, 2018). Therefore, tea can provide an excellent framework in machine learning methodologies because of its ability to provide a large number of progenies to be tested or studied. Also, with the recent availability of tea genome data, genotype and phenotype data can be integrated and thus, can provide valuable genetic and genomic resources to plant breeders to uncover novel trait-associated candidate genes. The current study was carried out for the first time in tea using genome-enabled prediction models to predict black tea quality and drought tolerance traits with the aim of reducing cost and time in tea breeding due to its long selection cycle.

## Materials and methods

### Genome selection dataset

The genomic dataset used in this study comprises of 255 F_1_ individuals (discovery population) from two segregating tea populations (TRFK St. 504 and TRFK St. 524) and 56 individual tea cultivars (validation population) with each individual phenotyped for black tea quality traits and drought tolerance trait. The selected eight phenotypic traits used were, %RWC, caffeine, catechin and tea tasters’ scores (aroma, astringency, brightness, briskness, and colour) which are associated with black tea quality and drought tolerance traits. All the individuals in both the discovery and validation populations were genotyped using 1 421 DArTseq markers (Koech *et al.*, 2018; Koech *et al.*, 2019).

### Transforming and ranking phenotypic data

The phenotypic data (*y*) in 311 tea datasets were centred at zero and standardised to unit variance. The *y* in 310 datasets were then grouped into three classes (upper, middle and lower), based on 15-85 % percentiles of each phenotypic trait analysed. For example, for 15-85 % percentiles, 0.15 and 0.85 quantiles were used to split *y* into three classes: upper class, if *y* > 0.85; middle class, if 0.15 < *y* ≤ 0.85; and lower class; if *y* ≤ 0.15.

### Partition designs and cross-validation of the dataset

The analyses were carried out using R v 3.5.0 (2018). The caret package (Williams *et al.*, 2018) was the main package used for training and fitting the various models. Two-fold cross-validation was used for training the models on the discovery population, while the “best” models were consequently fitted to the validation population. The discovery population consisted of all the observations labelled “Discovery”, except for two parents (GW Ejulu L and TRFK 303/577) that were added to the validation set. The markers were averaged across the putative QTLs, annotated proteins and the KEGG pathways. For example, if, “pathway A” “QTL A” or “protein A” is expressed in ten out of the 13 000 DArTseq markers consisting of 0 and 1 score values, i.e. absence or presence of polymorphism in the genomic representation of the sample, then the ten, 0 and 1 score values are summed and divided by ten, which is the average of ten DArTseq markers. If only nine DArTseq markers are present in the data, then the 0 and 1 score values are summed and divided by nine, which is the average of nine DArTseq markers. The missing values in the dataset were imputed with 0 (i.e. not-expressed). The trained prediction models obtained were then applied to predict phenotypic trait values of individuals from the testing set labelled “Validation” using genomic, phenotypic, putative QTLs, annotated proteins and KEGG pathways data.

### Prediction models

The prediction models were built for the various phenotypes as dependent variables which resulted in four models namely: putative QTLs + annotated protein + KEGG pathway, putative QTLs, annotated protein and KEGG pathway. In all cases, the aggregated markers were used as independent variables (inputs to the models).

### Comparison of performance of different prediction models

The prediction models were ranked according to the root-mean-squared-error (RMSE) values, from smallest to largest. The variable of importance was calculated as a percentage between 0 and 100, depending on the model used from the varImp function (Williams *et al.*, 2018). The average variable of importance for each variable across all the prediction models fitted were calculated and weighted by the RMSE value for that particular model. The reason was to minimise overfitting the different prediction models. The percentage concordance was calculated by creating a binary version of each phenotype: high > = 0, and low = otherwise.

## Results

### Percent RWC

The best prediction models based on QTLs + annotated proteins + KEGG pathway for %RWC trait in both discovery and validation datasets based on ranked RMSE and percent concordance were the Principal Component Analysis, Supervised Principal Component Analysis and Independent Component Regression (Table 1). The percent concordance for all the three prediction models for the discovery population was 56.9%, and the RMSE ranged between 0.995 and 0.988. However, the two best prediction models for the validation population were Principal Component Analysis and Supervised Principal Component Analysis with RMSE of 0.986 and 0.990, respectively and 60.3% concordance. Principal Component Analysis and Supervised Principal Component Analysis models had better prediction ability in the validation population as compared to discovery population.

**Table 1:**
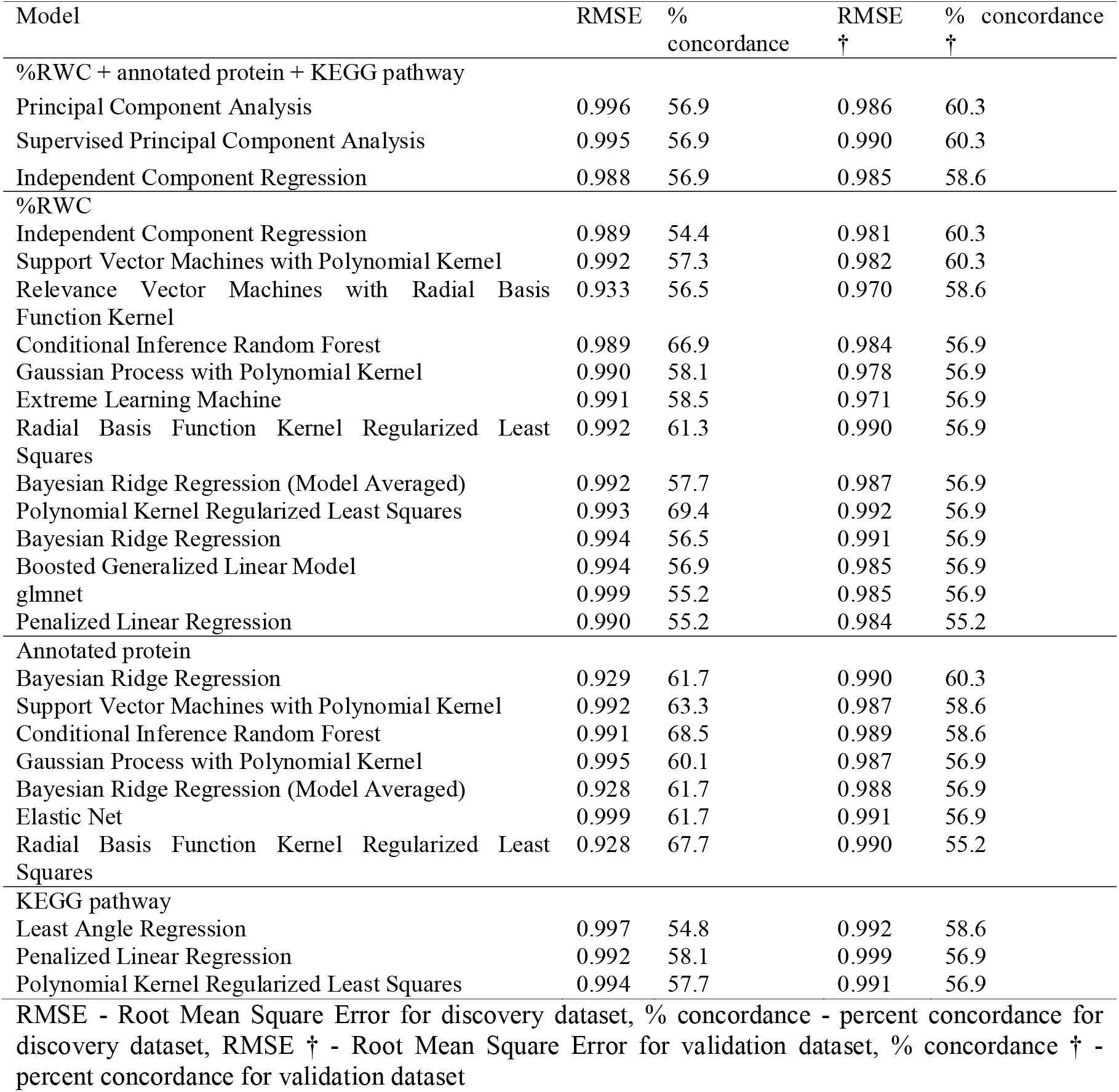
The best prediction models based on putative QTLs for %RWC trait, annotated protein and KEGG pathway in both discovery and validation datasets

The prediction models for %RWC trait based only on putative QTLs were cross-validated in both the discovery and validation populations. The two best predictive models with high percent concordance in the discovery population were Polynomial Kernel Regularized Least Squares (69.4%) and Conditional Inference Random Forest (66.9%) with RMSE of 0.993 and 0.989, respectively (Table 1). However, in the validation population, both prediction models had a percent concordance of 56.9% and RMSE of 0.992 and 0.984, which was lower than prediction models in discovery population. The two prediction models that had better predicted ability in the validation population with a 60.3% concordance were Independent Component Regression and Support Vector Machines with Polynomial Kernel (Table 1). The RMSE for Independent Component Regression and Support Vector Machines with Polynomial Kernel prediction models was 0.981 and 0.982, respectively. However, the predictive ability of the two models was also lower in the discovery population with 54.4% and 57.3% concordance, respectively.

The prediction models that had better prediction ability based only on functional annotation of putative proteins identified for %RWC trait were cross-validated in both datasets. Conditional Inference Random Forest (68.5%) and Radial Basis Function Kernel Regularized Least Squares (67.7%) prediction models had better predictive ability in the discovery population while Bayesian Ridge Regression, Support Vector Machines with Polynomial Kernel and Conditional Inference Random Forest models had better predictive ability of 60.3%, 58.6%, and 58.6%, respectively in the validation population (Table 1). The RMSE for prediction models that had better prediction ability in both dataset ranged between 0.928 and 0.991. Conditional Inference Random Forest was the only model had better prediction ability in both the discovery and validation populations. The models that performed better in prediction of putative QTLs based only on KEGG pathway in both datasets were Least Angle Regression, Penalized Linear Regression and Polynomial Kernel Regularized Least Squares (Table 1). The percent concordance for both datasets ranged between 54.8% and 58.6% with RMSE of between 0.991 and 0.999.

### Caffeine

The four prediction models based on QTLs + annotated proteins + KEGG pathway for caffeine trait that performed better in the discovery dataset were the Tree-Based Ensembles (61.5%), Linear Regression with Forward Selection (59.1%), Tree Models from Genetic Algorithms (57.5%) and Least Angle Regression (57.5%). Also, the four models performed better in the validation dataset with a 65% concordance for each model (Table 2).

**Table 2:**
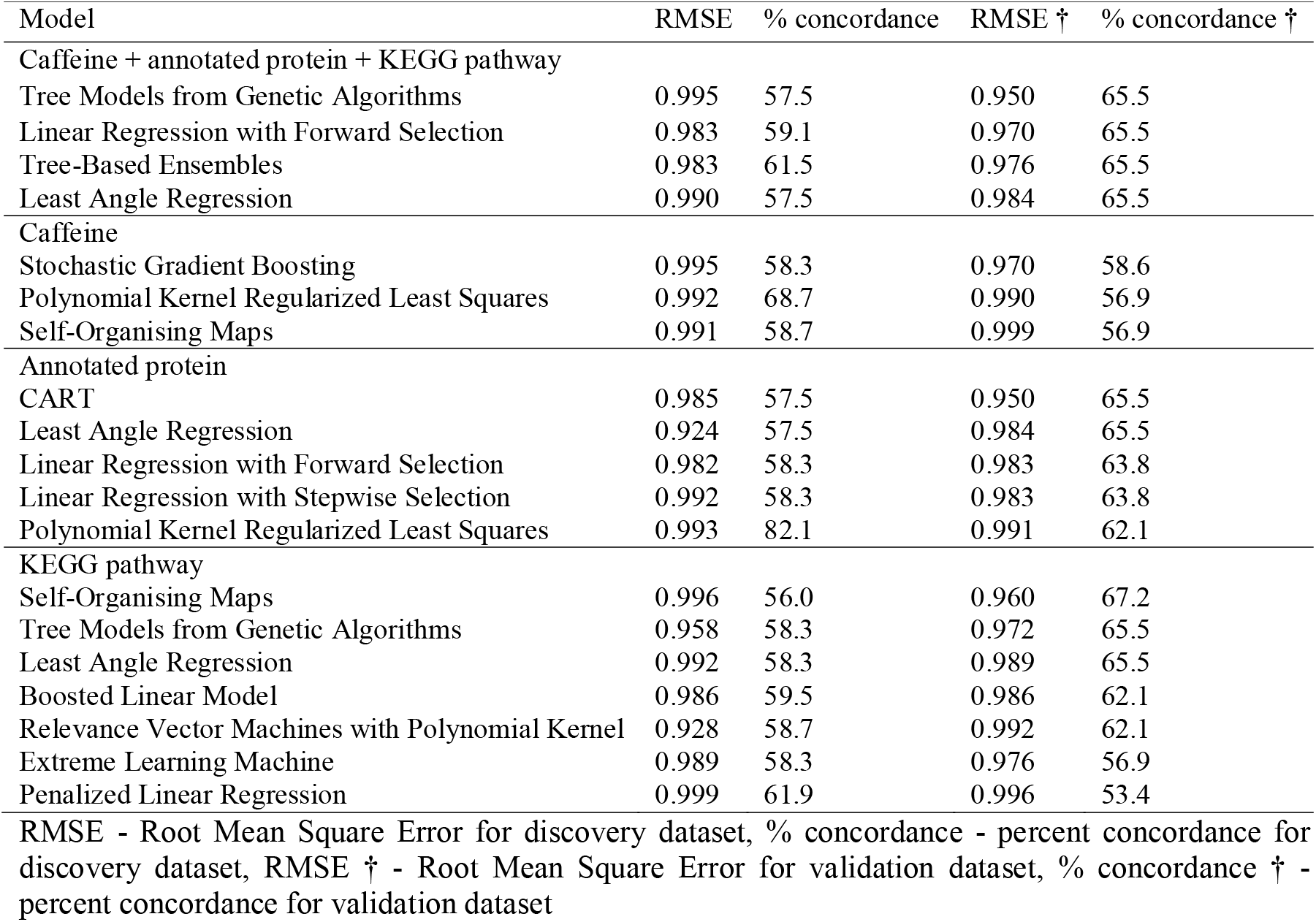
The best prediction models based on putative QTLs identified for caffeine trait, annotated protein and KEGG pathway in both discovery and validation dataset

The best prediction model based on only putative QTLs for caffeine trait in both the discovery and validation datasets were Polynomial Kernel Regularized Least Squares, Self-Organising Maps, and Stochastic Gradient Boosting. Polynomial Kernel Regularized Least Squares had a high percent concordance of 68.7% in the discovery dataset and 56.9% in the validation dataset. Stochastic Gradient Boosting performed better in the validation dataset with percent concordance of 58.6% than in the discovery dataset which had percent concordance of 58.3%.

The prediction models for caffeine trait based only on annotated proteins were cross-validated in both the discovery and validation datasets. A total of five models which performed better in both discovery and validation datasets (Table 2). The Polynomial Kernel Regularized Least Squares model had a high percent concordance of 82.1% and 62.1% in discovery and validation dataset, respectively. However, the best prediction models in validation dataset were Classification and Regression Tree (CART) (65.5%), and Least Angle Regression (65.5%) which both had a percent concordance of 57.5% in the discovery dataset. A total of seven KEGG pathway-based prediction models for caffeine trait performed better in both discovery and validation datasets. The percent concordance for the discovery dataset, ranged between 56% and 61.9% for Self-Organising Maps and Penalized Linear Regression, respectively. However, for the validation dataset, Penalized Linear Regression had percent concordance of 53.4% and 67.2% for Self-Organising Maps (Table 2). The RMSE for both discovery and validation dataset ranged between 0.958 and 0.999.

### Catechin

Similarly, the prediction models based on QTLs + annotated proteins + KEGG pathway for catechin trait were built and cross-validated in the discovery and validation datasets. Out of the 98 build prediction models, Extreme Learning Machine was the only prediction model that had low RMSE (0.942 and 0.978) and had better predictive ability for catechin trait in both the discovery and validation datasets. The percent concordance in the discovery and validation datasets was 55.6% and 69%, respectively (Table 3).

**Table 3:** The best prediction model based on putative QTLs identified for catechin trait, annotated protein and KEGG pathway in both discovery and validation dataset

| Model | RMSE | % concordance | RMSE † | % concordance † |
| --- | --- | --- | --- | --- |
| Catechin + annotated protein + KEGG pathway |  |  |  |  |
| Extreme Learning Machine | 0.942 | 55.6 | 0.978 | 69.0 |
| Catechin |  |  |  |  |
| Principal Component Analysis | 0.924 | 68.7 | 0.991 | 65.5 |
| Annotated protein |  |  |  |  |
| Neural Network | 1.22 | 55.6 | 1.12 | 65.5 |
| KEGG pathway |  |  |  |  |
| Self-Organising Maps | 0.972 | 54.0 | 0.969 | 75.9 |
| Independent Component Regression | 0.974 | 54.4 | 0.994 | 69.0 |

RMSE - Root Mean Square Error for discovery dataset, % concordance - percent concordance for discovery dataset, RMSE † - Root Mean Square Error for validation dataset, % concordance † - percent concordance for validation dataset

Also, Principal Component Analysis was the only best prediction model based on putative QTLs identified for catechin trait in both the discovery and validation dataset (Table 3). The prediction ability of Principal Component Analysis model was 68% with an RMSE of 0.924 in the discovery dataset and 65.5% in the validation dataset with an RMSE of 0.991.

The prediction models for catechin trait based on only annotated QTLs were cross-validated in both the discovery and validation datasets. Neural Network was the only prediction model that had better prediction ability in both the discovery and validation datasets with 55.6% and 65.6% concordance, respectively (Table 3). However, the prediction model in both discovery and validation dataset had a high RMSE value of 1.22.

The two prediction models that had high prediction ability for catechin trait based on only KEGG pathway in both the discovery and validation datasets were Self-Organising Maps and Independent Component Regression prediction models (Table 3). In the discovery dataset, Self-Organising Maps model had 54% concordance with RMSE of 0.972 while Independent Component Regression model had 54.4% concordance with RMSE of 0.974. However, in the validation dataset, Self-Organising Maps and Independent Component Regression prediction models had high percent concordance of 75.9% and 69% but with high RMSE of 0.969 and 0.994, respectively (Table 3).

### Aroma

Bayesian Ridge Regression (63.8%), The Bayesian LASSO (61.9%) and Least Angle Regression (59%) prediction models had high prediction ability for aroma based on QTLs + annotated proteins + KEGG pathway approach in the discovery dataset. The three prediction models also had high prediction ability in the validation dataset but with 62.7% concordance for Least Angle Regression model (Table 4).

**Table 4:** The best prediction model based on putative QTLs identified for tea liquor aroma trait, annotated protein and KEGG pathway in both discovery and validation dataset

| Model | RMSE | % concordance | RMSE † | % concordance † |
| --- | --- | --- | --- | --- |
| Aroma + annotated protein + KEGG pathway |  |  |  |  |
| Least Angle Regression | 0.996 | 59.0 | 0.988 | 62.7 |
| The Bayesian LASSO | 0.964 | 61.9 | 0.986 | 56.9 |
| Bayesian Ridge Regression (Model Averaged) | 0.969 | 63.8 | 0.985 | 54.9 |
| Aroma |  |  |  |  |
| The Bayesian LASSO | 0.994 | 56.2 | 0.986 | 66.7 |
| Bayesian Ridge Regression (Model Averaged) | 0.994 | 61.0 | 0.986 | 66.7 |
| Relevance Vector Machines with Polynomial Kernel | 0.994 | 62.4 | 0.957 | 66.7 |
| Radial Basis Function Kernel Regularized Least Squares | 0.994 | 73.8 | 0.987 | 64.7 |
| Penalized Linear Regression | 0.990 | 59.5 | 0.966 | 64.7 |
| Polynomial Kernel Regularized Least Squares | 0.994 | 79.5 | 0.989 | 62.7 |
| Elasticnet | 0.989 | 59.0 | 0.979 | 62.7 |
| Least Angle Regression | 0.989 | 59.0 | 0.979 | 62.7 |
| Boosted Linear Model | 0.991 | 60.5 | 0.975 | 62.7 |
| Gaussian Process with Polynomial Kernel | 0.994 | 61.9 | 0.972 | 62.7 |
| Independent Component Regression | 1.000 | 50.5 | 0.971 | 62.7 |
| The LASSO | 0.990 | 59.0 | 0.971 | 62.7 |
| Least Angle Regression | 0.990 | 59.0 | 0.971 | 62.7 |
| Boosted Generalized Linear Model | 1.000 | 60.0 | 0.966 | 62.7 |
| Conditional Inference Tree | 0.994 | 59.0 | 0.965 | 62.7 |
| Multi-Step Adaptive MCP-Net | 0.948 | 59.0 | 0.960 | 62.7 |
| Ridge Regression with Variable Selection | 0.999 | 59.0 | 0.981 | 60.8 |
| Relaxed LASSO | 0.994 | 60.5 | 0.966 | 60.8 |
| Random Forest | 0.992 | 100.0 | 0.993 | 58.8 |
| Linear Regression with Backwards Selection | 0.992 | 59.0 | 0.981 | 58.8 |
| Linear Regression with Forward Selection | 0.992 | 59.0 | 0.981 | 58.8 |
| Linear Regression with Stepwise Selection | 0.992 | 59.0 | 0.981 | 58.8 |
| Tree-Based Ensembles | 0.978 | 74.8 | 0.986 | 54.9 |
| Annotated protein |  |  |  |  |
| Parallel Random Forest | 1.02 | 99.5 | 0.987 | 60.8 |
| Regularized Random Forest | 1.01 | 98.6 | 0.992 | 56.9 |
| KEGG pathway |  |  |  |  |
| Principal Component Analysis | 0.994 | 54.3 | 0.974 | 62.7 |
| Tree-Based Ensembles | 1.001 | 59.5 | 0.982 | 58.8 |

RMSE - Root Mean Square Error for discovery dataset, % concordance - percent concordance for discovery dataset, RMSE † - Root Mean Square Error for validation dataset, % concordance † - percent concordance for validation dataset

Twenty-two prediction models with high prediction ability based on only putative QTLs identified for aroma trait were cross-validated in both discovery and validation datasets. The percent concordance in the discovery dataset ranged between 56.2% and 100%. Random Forest and The Bayesian LASSO were the models with high predictive ability for aroma trait based on only putative QTLs identified approach (Table 4).

The Parallel Random Forest and Regularized Random Forest prediction models had high prediction ability for black tea aroma trait based on annotated proteins approach in both discovery and validation datasets (Table 4). The percent concordance for the two prediction models was 99.5% and 98.6%, respectively in the discovery dataset and 60.8% and 56.9%, respectively in the validation dataset. Although the discovery dataset had high percent concordance, the RMSE was higher than of the validation dataset.

Principal Component Analysis and Tree-Based Ensembles were prediction models that had high prediction ability for aroma trait based on only the KEGG pathway in both discovery and validation datasets (Table 4). In the discovery dataset, Principal Component Analysis and Tree-Based Ensembles prediction models had 54.3% and 59.5% concordance, respectively. The prediction ability of Principal Component Analysis model and Tree-Based Ensembles model in the validation dataset was 62.7% and 58.8%, respectively.

### Astringency

Extreme Learning Machine was the only prediction model with high prediction ability for black tea astringency trait based on QTLs + annotated proteins + KEGG pathway in both discovery and validation dataset with 56.2% and 60.8% concordance, respectively (Table 5). Random Forest, Tree-Based Ensembles, glmnet, and Least Angle Regression were prediction models with high prediction ability based on putative QTLs identified for black tea astringency trait with 93.3%, 70%, 60% and 59.5% concordance, respectively. However, in the validation dataset, percent concordance ranged between 54.9% and 56.9% with RMSE of between 1.004 and 1.054.

**Table 5:**
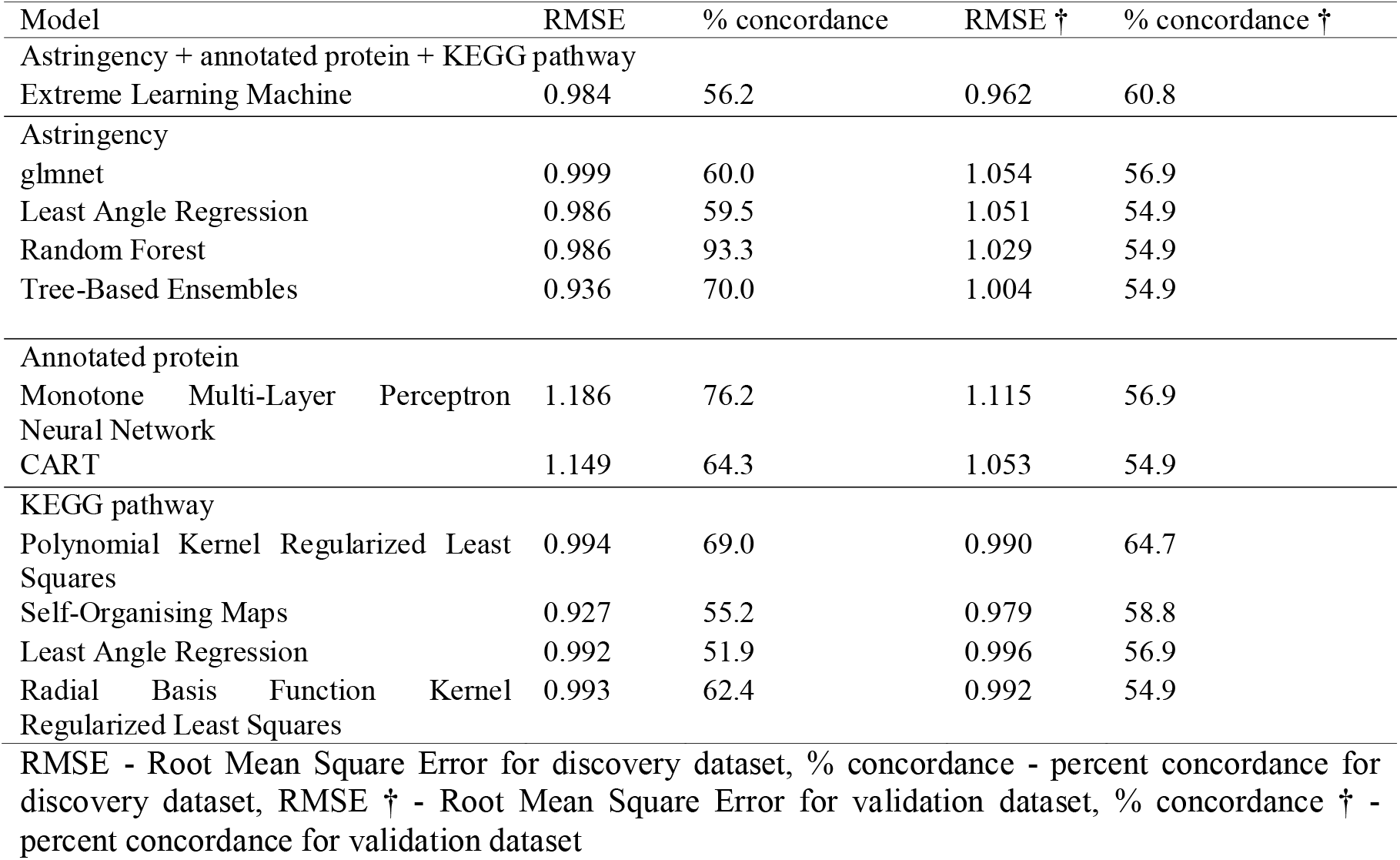
The best prediction model based on putative QTLs identified for tea liquor astringency trait, annotated protein and KEGG pathway in both discovery and validation dataset

The prediction models with high prediction ability in both datasets based on annotated proteins for putative QTLs identified for black tea astringency were Monotone Multi-Layer Perceptron Neural Network and CART (Table 5). Monotone Multi-Layer Perceptron Neural Network and CART had 76.2% and 64.3% concordance, respectively in the discovery dataset, whereas in the validation dataset, the two prediction models had 56.9% and 54.9% concordance, respectively. However, the two models had high RMSE in both the discovery and validation datasets. On KEGG pathway-based prediction models for putative QTLs identified for black tea astringency traits, Polynomial Kernel Regularized Least Squares model had high percent concordance of 69% and 64.7% in the discovery and validation dataset, respectively (Table 5).

### Brightness

All the four prediction models for black tea brightness trait based on QTLs + annotated proteins + KEGG pathway in the discovery dataset had low percent concordance of 51% with genomic DArTseq markers (Table 6) However, in the validation dataset, all the four prediction models had high percent concordance with genomic DArTseq markers. The prediction ability of Extreme Learning Machine model was 71.2% while Non-Informative Model, partDSA and Stacked AutoEncoder Deep Neural Network model was each 69.5%, respectively (Table 6).

**Table 6:** The best prediction model based on putative QTLs identified for tea liquor brightness trait, annotated protein and KEGG pathway in both discovery and validation dataset

| Model | RMSE | % concordance | RMSE † | % concordance † |
| --- | --- | --- | --- | --- |
| Brightness + annotated protein + KEGG pathway |  |  |  |  |
| Extreme Learning Machine | 0.974 | 51.9 | 0.966 | 71.2 |
| Non-Informative Model | 0.998 | 51.0 | 0.990 | 69.2 |
| partDSA | 0.998 | 51.0 | 0.990 | 69.2 |
| Stacked AutoEncoder Deep Neural Network | 0.994 | 51.0 | 0.999 | 69.2 |
| Brightness |  |  |  |  |
| Non-Informative Model | 0.998 | 51.0 | 0.990 | 69.2 |
| CART | 0.985 | 51.0 | 0.990 | 69.2 |
| Annotated protein |  |  |  |  |
| Polynomial Kernel Regularized Least Squares | 0.997 | 79.0 | 0.990 | 53.8 |
| KEGG pathway |  |  |  |  |
| Supervised Principal Component Analysis | 0.991 | 56.2 | 0.993 | 59.6 |

RMSE - Root Mean Square Error for discovery dataset, % concordance - percent concordance for discovery dataset, RMSE † - Root Mean Square Error for validation dataset, % concordance † - percent concordance for validation dataset

The prediction models based on putative QTLs identified for black tea brightness trait in the discovery dataset also had low percent concordance with genomic DArTseq markers (Table 6). However, Polynomial Kernel Regularized Least Squares was the best prediction model in both datasets with 79% and 53.8% concordance in the discovery and validation dataset, respectively.

Supervised Principal Component Analysis model had high prediction ability in both discovery and validation dataset based on the KEGG pathway for putative QTLs identified for black tea brightness trait with 56.2% and 59.6%, respectively (Table 6).

### Briskness

A total of eight prediction models had high prediction ability for black tea liquor briskness trait based on QTLs + annotated proteins + KEGG pathway in both discovery and validation datasets (Table 7). Stacked AutoEncoder Deep Neural Network, Random Forest Rule-Based Model, Quantile Regression with LASSO penalty, Non-Convex Penalized Quantile Regression and Cubist had high prediction ability of 65.2% and 74% concordance in both discovery and validation datasets, respectively.

**Table 7:** The best prediction model based on putative QTLs identified for tea liquor briskness trait, annotated protein and KEGG pathway in both discovery and validation dataset

| Model | RMSE | % concordance | RMSE † | % concordance † |
| --- | --- | --- | --- | --- |
| Briskness + annotated protein + KEGG pathway |  |  |  |  |
| Stacked AutoEncoder Deep Neural Network | 0.994 | 65.2 | 0.995 | 74.0 |
| Random Forest Rule-Based Model | 0.997 | 65.2 | 0.992 | 74.0 |
| Quantile Regression with LASSO Penalty | 0.997 | 65.2 | 0.992 | 74.0 |
| Non-Convex Penalized Quantile Regression | 0.997 | 65.2 | 0.992 | 74.0 |
| Cubist | 0.997 | 65.2 | 0.992 | 74.0 |
| Linear Regression with Forward Selection | 0.999 | 57.6 | 0.979 | 66.0 |
| Linear Regression with Stepwise Selection | 0.999 | 57.6 | 0.979 | 66.0 |
| Extreme Learning Machine | 0.980 | 56.2 | 0.990 | 56.0 |
| Briskness |  |  |  |  |
| Quantile Regression with LASSO penalty | 0.997 | 65.2 | 0.992 | 74.0 |
| Non-Convex Penalized Quantile Regression | 0.997 | 65.2 | 0.992 | 74.0 |
| Cubist | 0.997 | 65.2 | 0.992 | 74.0 |
| Annotated protein |  |  |  |  |
| Quantile Regression with LASSO penalty | 0.997 | 65.2 | 0.992 | 74.0 |
| Non-Convex Penalized Quantile Regression | 0.997 | 65.2 | 0.992 | 74.0 |
| Polynomial Kernel Regularized Least Squares | 0.998 | 80.0 | 0.990 | 54.0 |
| KEGG pathway |  |  |  |  |
| Quantile Regression with LASSO penalty | 0.997 | 65.2 | 0.992 | 74.0 |
| Non-Convex Penalized Quantile Regression | 0.997 | 65.2 | 0.992 | 74.0 |
| Cubist | 0.997 | 65.2 | 0.992 | 74.0 |
| Multi-layer Perceptron Network by Stochastic Gradient Descent | 0.994 | 65.2 | 0.998 | 74.0 |
| Multi-Layer Perceptron | 0.994 | 65.2 | 0.990 | 74.0 |
| Multi-Layer Perceptron, multiple layers | 0.994 | 65.2 | 0.990 | 74.0 |
| Extreme Learning Machine | 0.991 | 65.2 | 0.990 | 74.0 |

RMSE - Root Mean Square Error for discovery dataset, % concordance - percent concordance for discovery dataset, RMSE † - Root Mean Square Error for validation dataset, % concordance † - percent concordance for validation dataset

Of all the prediction models, Quantile Regression with LASSO penalty, Non-Convex Penalized Quantile Regression and Cubist had high prediction ability based on only putative QTLs with 65.2% and 74% concordance in both discovery and validation datasets (Table 7). Similarly, Quantile Regression with LASSO penalty and Non-Convex Penalized Quantile Regression were better prediction models for black tea briskness based on annotated proteins (Table 7). There were seven prediction models with better prediction ability based on the KEGG pathway for putative QTLs identified for black tea liquor briskness trait with had high percent concordance of between 65.2% and 74% (Table 7).

### Colour

The prediction models for black tea liquor colour trait based on QTLs + annotated proteins + KEGG pathway were cross-validated in both the discovery and validation populations (Table 8). Polynomial Kernel Regularized Least Squares and Extreme Learning Machine were the only two models in both datasets that had high percent concordance with genomic DArTseq markers. The percent concordance of Polynomial Kernel Regularized Least Squares and Extreme Learning Machine prediction models were 55.7% and 54.3%, respectively in the discovery dataset, whereas, in the validation dataset, the two prediction models had 58.6% and 53.4% concordance, respectively.

**Table 8:**
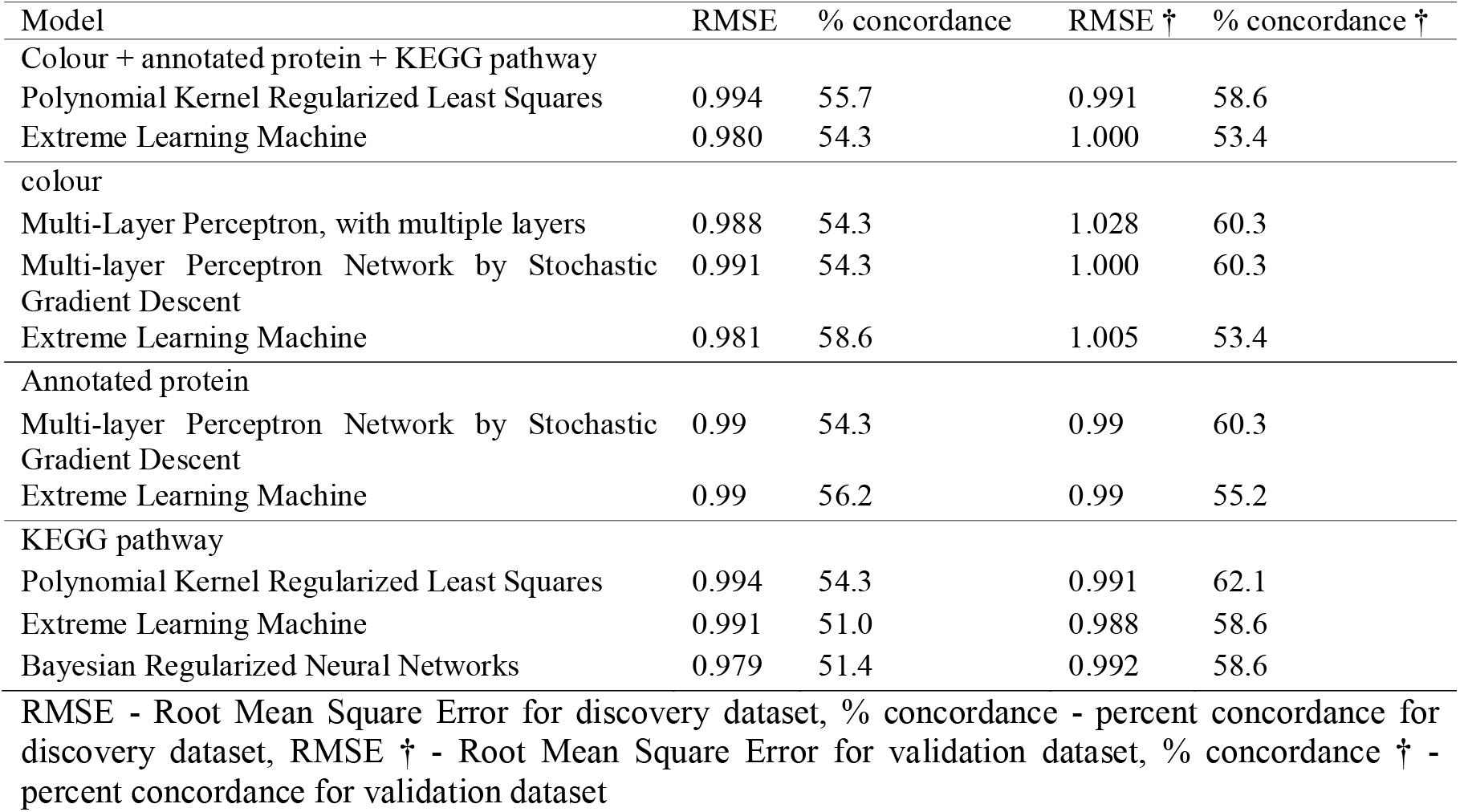
The best prediction model based on putative QTLs identified for tea liquor colour trait, annotated protein and KEGG pathway in both discovery and validation dataset

The prediction models that had better prediction ability in both datasets based only on putative QTLs identified for tea liquor colour trait were Multi-Layer Perceptron, with multiple layers and Multi-layer Perceptron Network by Stochastic Gradient Descent with 54.3% and 60.3% concordance, respectively (Table 8). The Multi-layer Perceptron Network by Stochastic Gradient Descent and Extreme Learning Machine models were better prediction models based on annotated proteins for black tea colour trait in both datasets (Table 8). In the discovery dataset, the Multi-layer Perceptron Network by Stochastic Gradient Descent model and the Extreme Learning Machine model were 54.3% and 56.2% concordance, respectively. Also, the two models had a better prediction ability in the validation dataset with 60.3% and 55.2% concordance, respectively. Polynomial Kernel Regularized Least Squares was the best prediction model based on KEGG pathway for putative QTLs identified for black tea liquor colour trait in both datasets (Table 8). The percentage concordance of Polynomial Kernel Regularized Least Squares model was 54.3% and 62.1% in the discovery and validation datasets, respectively.

The putatively annotated proteins for %RWC trait with the high percentage of importance were those associated with abiotic stress response while the metabolic KEGG pathways were those associated with arginine and proline metabolism (78.3%), glycerolipid metabolism (67.4%), and fructose and mannose metabolism (53.2%). The putatively annotated protein with high percentage of importance associated with caffeine trait in both discovery and validation datasets was N-(5’ phosphoribosyl) anthranilate (PRA) isomerase (91.9%). The metabolic KEGG pathways that are involved in carbon fixation in photosynthetic organisms (79.3%) and purine metabolism (63.7%) had high percentage of importance (Supplementary Table S1).

Phosphoribulokinase or uridine kinase family (45.1%) and autophagy-related protein 11 (32.9%) were variables of importance for annotated proteins associated with catechin trait in both discovery and validation dataset was alanine aspartate and glutamate metabolism (38.3%). Aminotransferase class I and II enzymes were the annotated protein with high percent variable of importance were associated with catechin (94.8%), astringency (81.6%), brightness (83.8%), briskness (86.3%) and colour (79.4%) phenotypic traits (Supplementary Table S1). Also, arginine biosynthesis was the variable of importance based on KEGG pathway associated with caffeine (60.5%), catechin (88.4%), astringency (78.3%), brightness (83.1%), briskness (83.2%), colour (79.6%) phenotypic traits. The annotated proteins with high percent variables of importance and associated with black tea aroma trait in both discovery and validation dataset was 2OG Fe II oxygenase superfamily with 42% concordance. The metabolic KEGG pathways with a high percentage of importance and were associated with black tea aroma trait were flavone and flavonol biosynthesis pathways with 52.8% concordance (Supplementary Table S1).

## Discussion

This study reports the first initiative of applying machine learning techniques to develop putative QTLs + annotated proteins + KEGG pathway, putative QTLs, annotated proteins and KEGG pathway-based prediction models for tea breeding. While previous research, particularly on tea breeding, has largely been focused on the conventional linkage and AM methods in selection of tea cultivars with desirable traits, no work has focused on prediction models to improve tea breeding. This study compared the performance of 98 different machine learning models which comprised of both supervised and unsupervised techniques. A few models were chosen as the best prediction models based on low RMSE and high percent concordance in both datasets. The low values of RMSE indicate high model quality (Alexander *et al.*, 2015; Lopatin *et al.*, 2016). Moreover, despite the rigorous and stringent validation procedures followed in this study for all the four based prediction approaches, putative QTLs + annotated proteins + KEGG pathway-based prediction approach, showed more robustness and usefulness in prediction than use of individual prediction based models. This important contribution also demonstrates the obvious underlying principle of the basic quantitative relationship between identified QTLs, functions of annotated proteins and the role of identified QTLs in the KEGG pathways.

In this study, different genomic prediction models were compared in terms of their predictive ability, or accuracy for different traits. However, the predictive ability of all models varied across traits. The difference in the prediction values of different prediction models for the same trait confirms that only traits that are easier to phenotype should be considered for genomic predictions (Nyine *et al.*, 2018). Similarly, the difference in model performance between traits suggests that variation in trait architecture, number of QTLs controlling the trait and LD between markers and QTL influence the performance of the models (Clark *et al.*, 2011). The results in this study are in agreement with previous studies in that prediction model that perform variable selection (possibly irrelevant variables) such as Extreme Learning Machine give better predictive ability values (Huang *et al.*, 2006). For example, Extreme Learning Machine model which is robust and sensitive to variable selection had better prediction ability for catechin, astringency, brightness, briskness, and colour based on putative QTLs + annotated proteins + KEGG pathway approach. The argument on the superiority of the model could also be highly dependent on the presence of large QTL effects. It is likely that the phenotypic traits that are controlled by large effect QTL in tea were selected by Extreme Learning Machine model in all cross-validations. The high level of concordance in the four prediction approaches also suggests that some molecular markers could be associated with the phenotypic traits in question. The same molecular markers are linked to proteins and metabolic pathways of different phenotypic traits. In addition, variable of importance for each trait based on either annotated proteins or KEGG pathway confirmed the predictive ability of each model.

The percent variable of importance of annotated proteins associated with %RWC trait were actin, armadillo beta-catenin-like repeat and isocitrate isopropyl malate dehydrogenase which are associated with abiotic stress (Drøbak *et al.*, 2004; Kamal *et al.*, 2012; Sharma *et al.*, 2014). The KEGG pathways were arginine and proline metabolism, glycerolipid metabolism and fructose and mannose metabolism, which are primary metabolites reported to be associated with abiotic stress (Upchurch, 2008; Keunen *et al.*, 2013; Majumdar *et al.*, 2016). The high percentage of importance of putatively annotated proteins for caffeine trait was N-(5’ phosphoribosyl) anthranilate (PRA) isomerase, which is involved in the first and intermediate pathway in the purine biosynthesis (D’Mello, 2017). Caffeine and other methylxanthines such as theobromine (3, 7-dimethylxanthine), methyluric acids and theophylline are present in tea and are classified as purine alkaloids (Ashihara *et al.*, 2008). Therefore, the putative protein, N-(5’ phosphoribosyl) anthranilate (PRA) isomerase may be associated with the biosynthesis of caffeine, theobromine, theophylline and methyluric acids in tea. Also, phenylalanine, tyrosine, and tryptophan are products of shikimate pathway which is involved in the biosynthesis of plant flavonoids, including catechins (Ghasemzadeh; Ghasemzadeh, 2011). For KEGG pathways, DArT markers with a high percentage of importance were those associated with carbon fixation in photosynthetic organisms, arginine biosynthesis and purine metabolism, which is associated with caffeine biosynthesis (Ashihara *et al.*, 2013). The putatively annotated proteins for catechin trait were those associated with phenylalanine, tyrosine and tryptophan biosynthesis and carbon fixation, which are products for shikimate pathway (Maeda;Dudareva, 2012) and abiotic stress response (stomatal closure). Hence, it was in agreement with this study since previous reports have shown that catechins play a role as indicators of desiccation tolerance in tea plants (Cheruiyot *et al.*, 2008). Aminotransferases have been shown to play a major role in a variety of metabolic pathways, including, amino acid biosynthesis and photorespiration. Aminotransferase I and II, putatively annotated for catechin trait, are involved in the biosynthesis of phenylalanine and tyrosine, which are precursors of flavonoid biosynthesis (Maeda *et al.*, 2011). Also, aminotransferase I and II had high percent variable of importance of annotated proteins associated with astringency, brightness, briskness and colour traits. In relation to this argument, theaflavins, and thearubigins present in black tea are formed from the oxidation of catechins during the manufacture of black tea. Besides caffeine and catechins, amino acids such as arginine contribute to astringency in tea infusion (Zhen, 2003). The findings corroborate with the current study on the variable of importance in the KEGG pathways which is involved in arginine biosynthesis.

In conclusion, the comparison of the performance of different prediction models was based on ranked RMSE values. The predictive ability of different machine learning methodologies applied were significantly different based on putative QTLs + annotated proteins + KEGG pathways, putative QTLs, annotated proteins and KEGG pathways approaches. However, the discovery and validation tea populations were phenotyped in different locations and in different years. Thus, the putative genotype by environment interactions due to differences in years and locations were not considered, and this might have affected the prediction accuracy. Conversely, since the %concordance in the validation population were quite high, this indicates that the models are robust across locations and years. The parents with their F_1_ progenies (discovery population) and the validation population were planted in the same plots, so there was an insufficient number of reference genotypes available to estimate genotype by environment interaction. Additional phenotyping of the discovery population in different locations is in progress, and using the average over multiple years may yield more stable estimates of the phenotypic performance, which could further increase the accuracy of the prediction models. Moreover, the results in this study suggest that for an outbreeding plant species such as tea, increasing the training population size could improve the predictive ability of the models as compared to using only two parental clones. In other tree species such as eucalyptus, pine and apples more than two parents have been used to improve on prediction ability and accuracy of the models. However, this research study has opened up a new avenue for future application of the machine learning techniques in tea breeding. Also, genomic selection can be applied earlier than phenotypic selection and thus reducing the time from initial crosses to planting in commercial tea fields. Implementation of genomic selection in an early-stage in tea breeding program can increase the selection intensity which is relevant concerning related traits which are often costly to phenotype.

## Acknowledgement

The authors acknowledge the financial support to conduct this research, and study grants for RK and PM from James Finlay (Kenya) Ltd., George Williamson (Kenya) Ltd., Sotik Tea Company (Kenya) Ltd., Mcleod Russell (Uganda) Ltd., the TRI of Kenya and Southern African Biochemistry and Informatics for Natural Products (SABINA). The *C. sinensis* cultivars used in this study were provided by the TRI of Kenya. Supplementary funding was provided by, the Technology and Human Resources for Industry Programme (THRIP), an initiative of the Department of Trade and Industries of South Africa (dti), the National Research Foundation (NRF) of South Africa, and the University of Pretoria (South Africa).

## Author Contribution

ZA, SK and RM were involved with the design of the experiment and plant material used. RK, PM and CN were involved in collection of plant material. RK performed the experiments. RK, PM, CN, SK, TL and ZA analyzed samples and interpreted the data. RK wrote the manuscript and revised by PM, CM, RM, SK, TLand ZA. The final manuscript was reviewed and approved by all the authors.

