## Supplementary Tables for "Genome-enabled prediction models for black tea (*Camellia sisnesnsis*) quality and drought tolerance traits"

**ELECTRONIC SUPPLEMENTARY MATERIALS**

The following information accompanies the article:

Supplementary Table 1. Percent variable of importance of annotated proteins and KEGG pathways related to black tea quality and drought tolerance traits

| %RWC | % variable of importance |
| --- | --- |
| Annotated protein | |
| Actin | 84.2 |
| Armadillo beta-catenin-like repeat | 78.8 |
| CSA002263 | 71.2 |
| Isocitrate isopropyl malate dehydrogenase | 66.3 |
| Peptidase C65 Otubain | 56.1 |
| BT1 family | 54.8 |
| Catalase | 48.9 |
| Autophagy-related protein 11 | 47.4 |
| KEGG pathway | |
| Arginine and proline metabolism | 78.3 |
| Glycerolipid metabolism | 67.4 |
| Alanine aspartate and glutamate metabolism | 55.8 |
| Tryptophan metabolism | 53.4 |
| Fructose and mannose metabolism | 53.2 |
| Caffeine | % variable of importance |
| Annotated protein | |
| N-(5’ phosphoribosyl anthranilate) (PRA) isomerase | 91.9 |
| Acyltransferase | 61.6 |
| Glutaminyl-tRNA synthetase | 60.6 |
| Glycosyl hydrolase family 9 | 41.8 |
| Peptidase C65 Otubain | 32.1 |
| KEGG pathway | |
| Carbon fixation in photosynthetic organisms | 79.3 |
| Purine metabolism | 63.7 |
| Arginine biosynthesis | 60.5 |
| Thiamine metabolism | 48.1 |
| Amino acid and nucleotide metabolism | 47.8 |
| Glyoxylate and dicarboxylate metabolism | 44.1 |
| Pyruvate metabolism | 30.9 |
| Cysteine and methionine metabolism | 30.1 |
| Catechin | % variable of importance |
| Annotated protein | |
| Diacylglycerol kinase catalytic domain | 105.6 |
| Aminotransferase class I and II | 94.8 |
| Phosphoribulokinase Uridine kinase family | 45.1 |
| CSA016461 | 34.4 |
| Autophagy-related protein 11 | 32.9 |
| CSA024230 | 31.5 |
| Thiolase C-terminal domain | 30.6 |
| KEGG pathway | |
| Arginine biosynthesis | 88.4 |
| Alanine aspartate and glutamate metabolism | 38.3 |
| Aroma | % variable of importance |
| Annotated protein | |
| Actin | 78.1 |
| CSA003424 | 49.0 |
| WD domain | 44.2 |
| 2OG Fe II oxygenase superfamily | 42.0 |
| 14-3-3 protein | 41.9 |
| NB-ARC domain | 40.3 |
| Pectinesterase | 39.2 |
| CSA033214 | 38.6 |
| N-(5’ phosphoribosyl) anthranilate (PRA) isomerase | 38.5 |
| ATPase family associated with various cellular activities AAA | 36.4 |
| KEGG pathway | |
| Tyrosine metabolism | 73.5 |
| Alanine aspartate and glutamate metabolism | 62.7 |
| Flavone and flavonol biosynthesis | 52.8 |
| Fructose and mannose metabolism | 48.6 |
| Pentose and glucuronate interconversions | 39.2 |
| Arginine biosynthesis | 38.5 |
| Astringency | % variable of importance |
| Annotated protein |  |
| Aminotransferase class I and II | 81.6 |
| Autophagy-related protein 11 | 42.5 |
| KEGG pathway | |
| Arginine biosynthesis | 78.3 |
| Arginine and proline metabolism | 71.1 |
| Glycin, serine and threonine metabolism | 51.0 |
| Brightness | % variable of importance |
| Annotated protein | |
| Aminotransferase class I and II | 83.8 |
| Autophagy-related protein 11 | 66.0 |
| Armadillo beta-catenin-like repeat | 65.6 |
| CSA024230 | 61.8 |
| CSA002263 | 61.1 |
| Phosphoribulokinase Uridine kinase family | 47.5 |
| Diacylglycerol kinase catalytic domain | 46.7 |
| ATPase family associated with various cellular activities AAA | 40.9 |
| KEGG pathway | |
| Arginine biosynthesis | 83.1 |
| Arginine and proline metabolism | 27.1 |
| Glycine, serine and threonine metabolism | 23.7 |
| Tyrosine metabolism | 21.1 |
| Briskness | % variable of importance |
| Annotated protein | |
| Aminotransferase class I and II | 86.3 |
| Armadillo beta-catenin-like repeat | 70.7 |
| CSA002263 | 66.0 |
| Autophagy-related protein 11 | 58.7 |
| CSA024230 | 57.0 |
| KEGG pathway | |
| Arginine biosynthesis | 83.2 |
| Arginine and proline metabolism | 33.7 |
| Terpenoid backbone biosynthesis | 29.0 |
| Phosphatidylinositol signalling system | 24.3 |
| Alanine aspartate and glutamate metabolism | 22.9 |
| Colour | % variable of importance |
| Annotated protein | |
| Aminotransferase class I and II | 79.4 |
| Phosphoribulokinase Uridine kinase family | 63.0 |
| Autophagy-related protein 11 | 54.6 |
| CSA024230 | 52.8 |
| Diacylglycerol kinase catalytic domain | 38.5 |
| Adaptor complexes medium subunit family | 31.0 |
| CSA016461 | 30.7 |
| ATPase family associated with various cellular activities AAA | 30.6 |
| Isocitrate isopropylmalate dehydrogenase | 26.9 |
| KEGG pathway | |
| Arginine biosynthesis | 79.6 |
| Porphyrin and chlorophyll metabolism | 46.9 |
| Arginine and proline metabolism | 41.7 |
| Pentose and glucuronate interconversions | 33.1 |
| Fructose and mannose metabolism | 28.6 |
| Glycine, serine and threonine metabolism | 27.9 |
